# Prediction of antiviral drugs against African Swine Fever Viruses based on protein-protein interaction analysis

**DOI:** 10.1101/599043

**Authors:** Zhaozhong Zhu, Yunshi Fan, Zena Cai, Zheng Zhang, Congyu Lu, Taijiao Jiang, Gaihua Zhang, Yousong Peng

**Affiliations:** College of Biology, Hunan University, Changsha, China; Suzhou Institute of Systems Medicine, Suzhou, China; Center of System Medicine, Institute of Basic Medical Sciences, Chinese Academy of Medical Sciences & Peking Union Medical College, Beijing, China; College of Life Sciences, Hunan Normal University, Changsha 410081, China

## Abstract

The African swine fever virus (ASFV) has severely influenced the swine industry of the world. Unfortunately, there is no effective antiviral drug or vaccine against the virus until now. Identification of new anti-ASFV drugs is urgently needed. Here, an up-to-date set of protein-protein interactions (PPIs) between ASFV and swine were curated by integration of PPIs from multiple sources. Thirty-two swine proteins were observed to interact with ASFVs and were defined as AIPs. They were found to play a central role in the swine PPI network, with significant larger degree, betweenness and smaller shortest path length than other swine proteins. Some of AIPs also interacted with several other viruses and could be taken as potential targets of drugs for broad-spectrum effect, such as HSP90AB1. Finally, the antiviral drugs which targeted AIPs and ASFV proteins were predicted. Several drugs with either broad-spectrum effect or high specificity on AIPs were identified, such as Polaprezinc. This work could not only deepen our understanding towards the ASFV-swine interactions, but also help for the development of effective antiviral drugs against the ASFVs.

## Introduction

African swine fever virus (ASFV), the causative agent of African swine fever (ASF), is an enveloped virus with double-stranded DNA of 170-193 kb. ASFV mainly infect suids and soft ticks. In swine populations, the virus can cause 100% mortality and severely influence the swine industry. The ASFV has caused ASF outbreaks in more than fifty countries in Africa, Europe, Asia, and South America until now ^1^. The latest reports showed that the virus has caused outbreaks in more than half of provinces in China ^2, 3^. How to effectively control the virus is still a great challenge for the globe ^4^.

Vaccine and antiviral drugs are believed to be the best tool for prevent viral infection and spread ^5^. Unfortunately, all the attempts to develop effective vaccines against ASFVs had failed. Therefore, it is in great need to develop effective antiviral drugs against ASFVs. Several studies have identified multiple compounds which could inhibit ASFV infections ^4^. For example, a recent study by Hakobyan et al. found that the rigid amphipathic fusion inhibitors displayed a potent and dose-dependent inhibitory effect on ASFV infection ^6^. Another recent study by Arabyan et al. showed that the genistein could hamper the ASFV infection by inhibiting the ASFV type II topoisomerase ^7^. However, all the antiviral drugs mentioned above have not been taken forward for commercial production. More candidate drugs are needed for further development.

Although most antiviral drugs target the viral proteins, in recent years several studies have attempted to develop antiviral drugs which targeted the host proteins. Compared to the drugs which targeted viral proteins, the drugs targeting host proteins had much more targets in the host cell ^8, 9^. Besides, they may be more mutant-insensitive since the host protein evolves much slower than viral proteins. With the rapid development of high-throughput assays, a large amount of protein-protein interactions (PPIs) between virus and host has been accumulated. Analysis of these PPIs in the perspective of network can help identify host proteins of importance for viral infection, which could be taken as potential targets for antiviral research ^8, 10^. For example, Han et al. predicted several antiviral drugs against human enterovirus 71 by systematic identification and analysis of PPIs between the virus and the host, suggesting the important role of PPI analysis on developing antiviral drugs targeting host proteins ^11^.

Here, we firstly curated a set of PPIs between ASFV and swine proteins by integration of PPIs from multiple sources; then, the swine proteins related to ASFV infection were identified; their roles in swine PPI network and in interacting with other viruses, and their functions were further investigated; finally, the candidate antiviral drugs targeting these swine proteins and ASFV proteins were predicted. This work could not only deepen our understanding towards the ASFV-swine interactions, but also help for the development of effective antiviral drugs against the ASFVs.

## Materials and Methods

### PPIs between ASFV and swine proteins

The PPIs between ASFV and swine proteins were compiled from three sources (Supplementary Table S1). First of all, 24 PPIs with median confidence (scores greater than 0.4) between ASFV and swine, were obtained from the database of Viruses.STRING ^12^ on January 8, 2019.

Secondly, 17 PPIs between ASFV and swine were obtained from the literature. This was achieved by firstly searching the PubMed database by the key word “ASFV” in the title or abstract on December 29, 2018, which resulted in 630 abstracts. Then, each abstract was manually screened based on whether it contained PPIs between ASFV and swine, and 117 abstracts were retained. Finally, the full texts of the manuscripts corresponding to these abstracts were read carefully and 17 extra PPIs between ASFV and swine were compiled from these papers.

Thirdly, 3 PPIs between ASFV and swine proteins were inferred based on sequence homology. This was conducted by firstly collecting viral proteins (except the ASFV) which interacted with swine proteins based on the database of Viruses.STRING. Then, 159 ASFV proteins encoded by BA71V, which were downloaded from NCBI RefSeq database ^13^, were blast ^14^ against these viral proteins. The hits with e-value smaller than 0.001, coverage greater than 40%, and sequence identity greater than 30%, were considered as homologs of ASFV proteins. The swine proteins which interacted with the hits were predicted to interact with the ASFV proteins.

### Swine PPI network

All the swine PPIs were downloaded from STRING database ^15^ on January 8, 2019. Only the PPIs with a median confidence (score greater than 0.4) were kept. Besides, the redundant PPIs were removed. Finally, a PPI network which consisted of 731,174 non-redundant PPIs between 18,683 swine proteins was obtained for further analysis.

### Network analysis and visualization

The igraph package (version 1.2.2)^16^ in R was used to analyze the topology of the PPI network. The degree and betweenness of proteins in the PPI network were calculated with the functions of *degree()* and *betweenness()*, respectively. The shortest path length between two proteins in the PPI network was calculated with the function of *shortest.paths()*.

The network was visualized with the help of Cytoscape (version 3.7.1)^17^.

### Functional enrichment analysis

The Gene Ontology (GO) terms and KEGG pathway enrichment analysis for the ASFV-interacting proteins (AIP) or the ASFV-associated proteins (AAP) were conducted with functions of *enrichGO()* and *enrichKEGG()* in the package “clusterProfiler” (version 3.6.0) ^18^ in R (version 3.4.2). All the GO terms and KEGG pathways with adjusted p-values smaller than 0.01 were considered as significant enrichment (Table S2).

### Prediction of candidate drugs targeting ASFV and swine proteins

Candidate drugs were predicted with the help of DrugBank (version 5.1.2) ^19^. The protein sequence of each ASFV protein encoded by BA71V, and that of each AIP was queried against DrugBank for similar targets with the default parameters. The drugs targeting the best hit were considered to be candidate drugs for the query protein. The properties of drugs, such as the type and group of drug, and ATC code, were also obtained from DrugBank (Supplementary Table S3).

## Results

### Interactions between ASFV and swine proteins

We firstly attempted to collect the interactions between ASFV and swine proteins as more as possible. In total, we obtained 44 protein-protein interactions (PPIs) between them (Figure 1A), including 24 PPIs from the database of Viruses.STRING, 17 PPIs from the literature and 3 PPIs inferred from PPIs between other virus and swines based on sequence homology (details in Materials and Methods). 16 ASFV proteins were involved in the PPIs. Half of ASFV proteins interacted with only one swine protein. For the remaining half of ASFV proteins, the DNA-directed DNA polymerase (DPOL) were found to interact with 13 swine proteins, while the Thymidine kinase (TDK) were found to interact with 6 swine proteins. 35 swine proteins were involved in the PPIs between ASFV and swines, which were defined as ASFV-interacting swine Proteins (AIPs). All of them only interacted with one ASFV protein except the proteins of DNAJA3, FBXO2 and SNAPIN.

**Figure 1.**
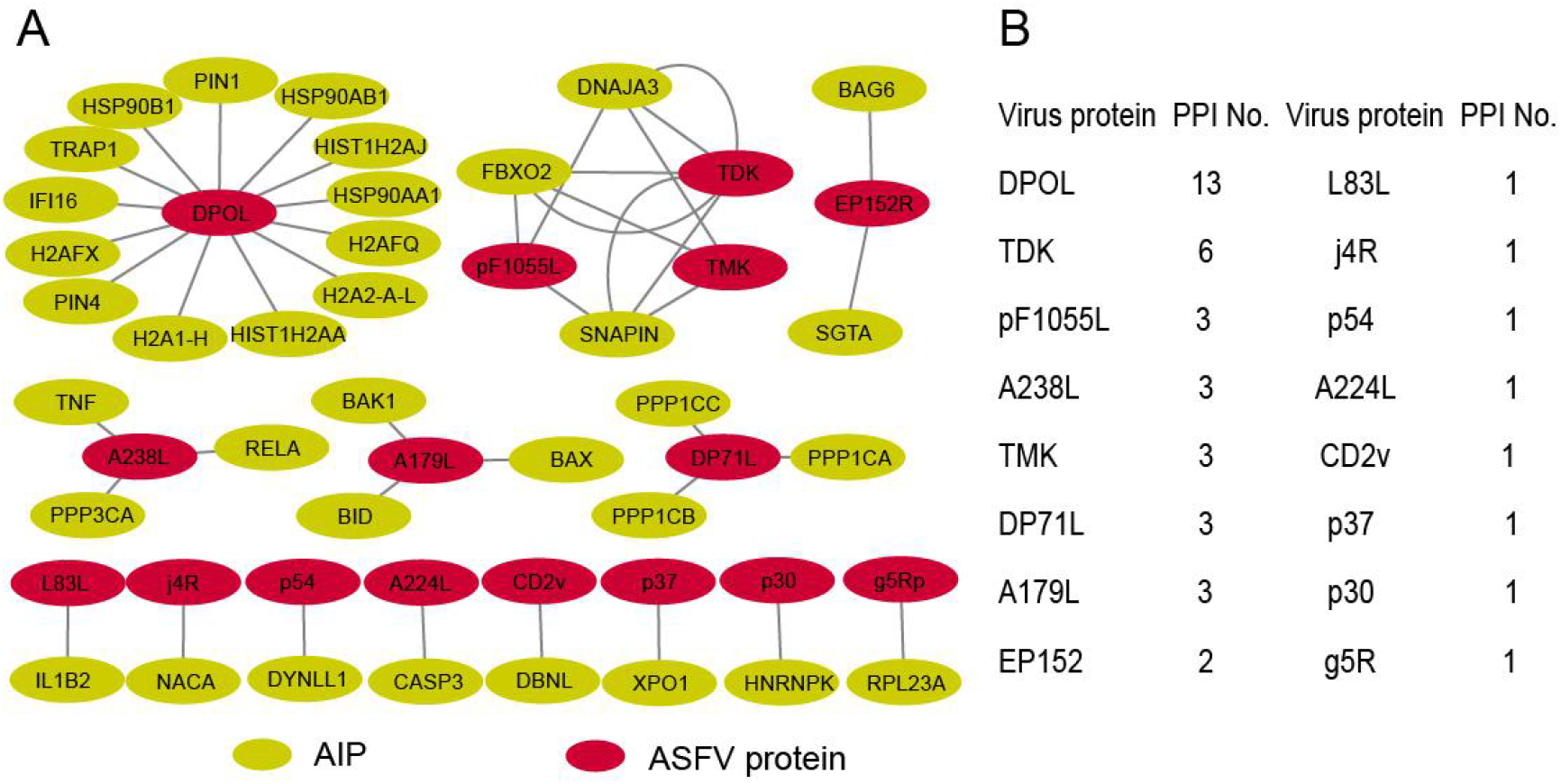
Overview of PPIs between the ASFV and swine. (A) Collected PPIs between ASFV and swine proteins; (B) All the ASFV proteins involved in PPIs and the number of interacted swine proteins.

### Construction of the ASFV-swine protein interaction network and topological analysis

To investigate the role of AIPs in the swine, a swine protein-protein interaction (PPI) network was constructed from the STRING database, which contained 731,174 non-redundant PPIs between 18,683 swine proteins. 32 (94%) AIPs were found to interact with other swine proteins. These proteins together with another 4028 swine proteins formed a protein interaction network with 8959 non-redundant interactions (Figure 2A), including 62 interactions between AIPs. We defined the 4028 AIP-interacting swine proteins as ASFV infection-associated swine proteins (AAPs).

**Figure 2.**
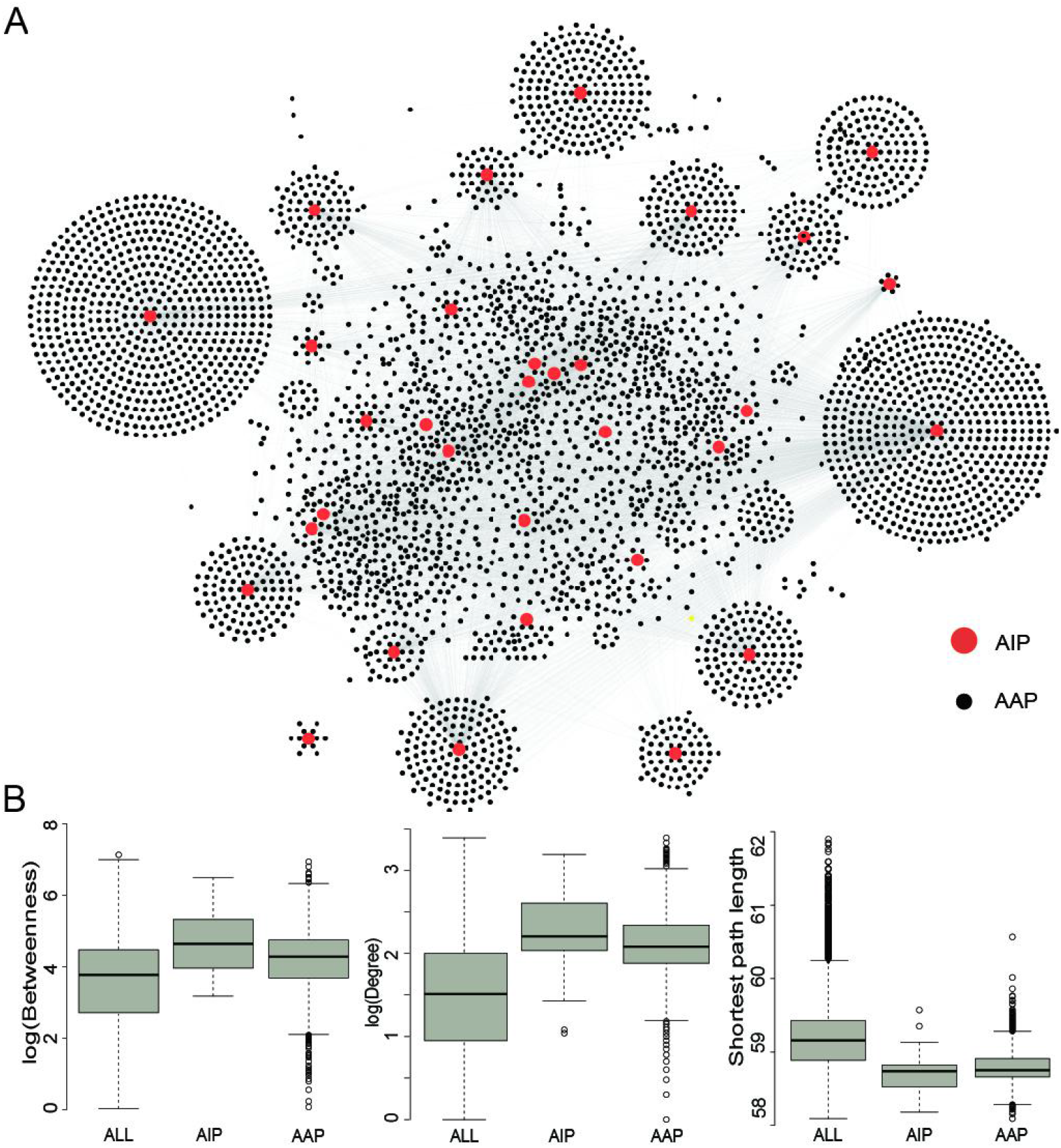
The AIP interaction network and the topological analysis of AIPs in the swine PPI network. (A) The PPI network between the AIPs (in red) and other swine proteins (in black). (B) Distribution of the degree, betweenness centrality and shortest path length for all proteins, AIPs and AAPs.

To investigate the centrality of AIPs in the swine PPI network, we calculated the degree and betweenness centrality, and the average shortest path length of each protein in the swine PPI network, including the AIPs and AAPs (Figure 2B). The median degree and betweenness centrality, and the median shortest path length of all proteins in the swine PPI network were 32, 59.2 and 10^3.5^, respectively, whereas these values for the AIPs were 161, 58.7 and 10^4.6^, respectively, and they were 121, 58.8 and 10^4^, respectively, for the AAPs (Figure 2B). The AIPs and AAPs were observed to have significant larger degrees and betweenness, and smaller shortest path length than all swine proteins, with p-values much smaller than 0.001 in the two-sided Wilcoxon rank-sum test. This suggested that the AIPs and AAPs played a central role in the swine PPI network.

### Functional analysis of AIPs and AAPs

Since the AIPs and AAPs were observed to play a central role in swine PPI network, we next investigated their functions. Functional enrichment analysis was conducted on the AIPs and AAPs (Table S2). Only a few GO terms in the domain of Molecular Function were enriched. Interestingly, more than 50 KEGG pathways were enriched in the AIPs (Table S2). The pathways of Necroptosis and Alcoholism were two of the most enriched pathways, both of which included more than 30% of all AIPs. Besides, three pathways related to virus infection, such as Human cytomegalovirus infection and Herpes simplex infection, were also enriched.

We further conducted the functional enrichment analysis on the AAPs. Figure 3 showed the top ten GO terms in three domains of GO and KEGG pathways enriched in the AAPs. In the domain of Biological Process, six of top ten enriched GO terms were related to cell death or apoptotic process; in the domain of Cellular Component, the AAPs were enriched in the nuclear and cytoskeleton; in the domain of Molecular Function, the AAPs were enriched in the GO terms of binding and enzyme activity. For the KEGG pathways, several signaling pathways were most enriched, such as PI3K-Akt signaling pathway and MAPK signaling pathway. Besides, several pathways related to virus infection were also enriched, such as the Human T-cell leukemia virus 1 infection.

**Figure 3.**
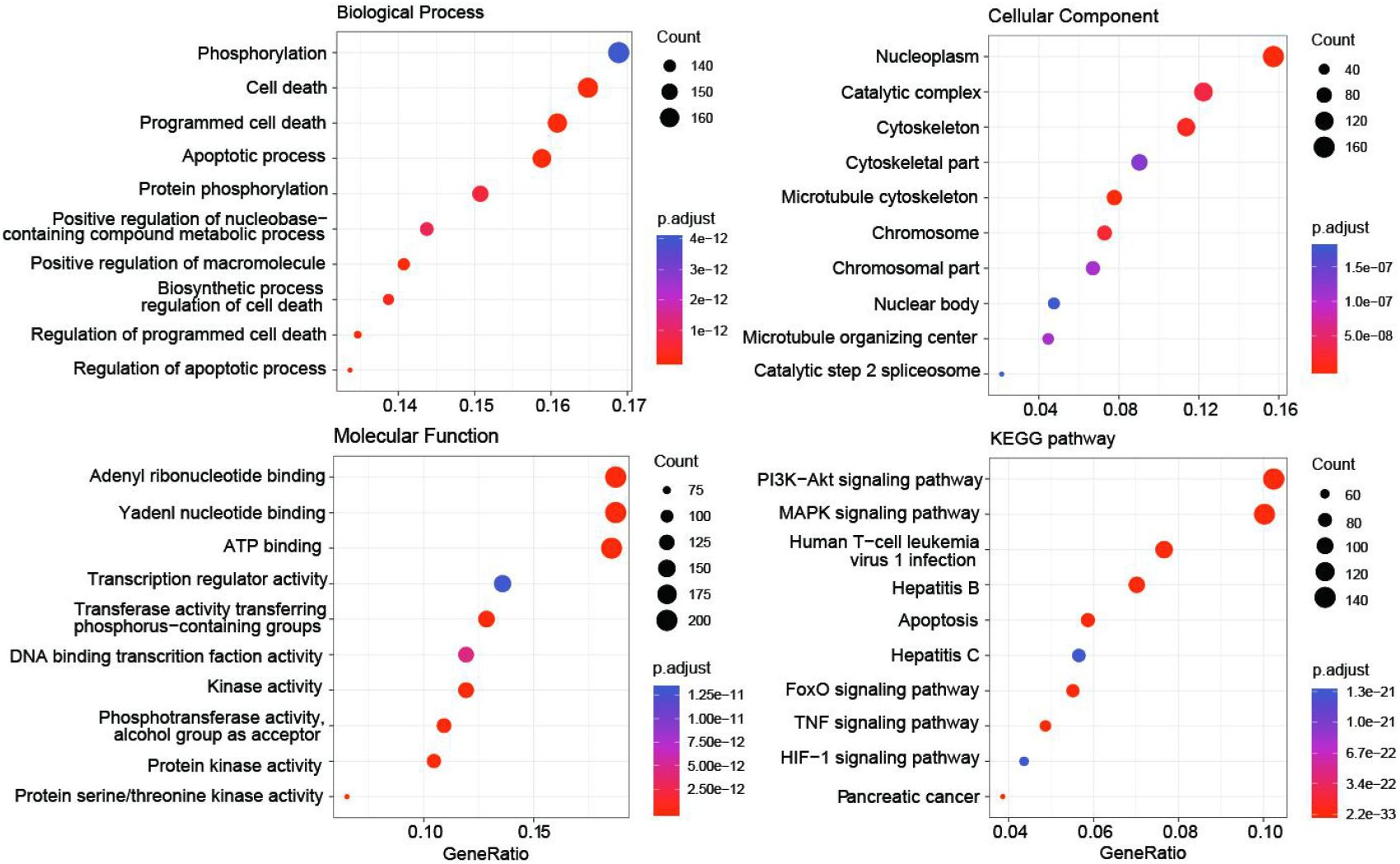
Functional enrichment analysis of AASPs. Top ten enriched terms in the domain of Biological Process, Cellular Component and Molecular Function, and KEGG pathways were shown.

### Overlap analysis of AIPs and other virus-interacting swine proteins

We then investigated the role of AIPs in the PPIs between swine and other viruses. All the PPIs between AIPs and other viral proteins which were public available in the Viruses.STRING database were obtained. As was shown in Figure 4A, 48 PPIs were obtained and shaped a network, which included 15 proteins of 11 other viruses (nodes in square) and 15 AIPs. 12 AIPs were observed to interact with more than one other virus. Besides, they also interacted with several hundreds of proteins in the swine PPI network (Figure 4B). For example, the heat shock protein 90s, including HSP90AB1, HSP90AA1 and HSP90B1, could interact with proteins from five other viruses, and interact with 1602 swine proteins, suggesting their central roles in both the virus-swine PPI network and swine PPI network.

**Figure 4.**
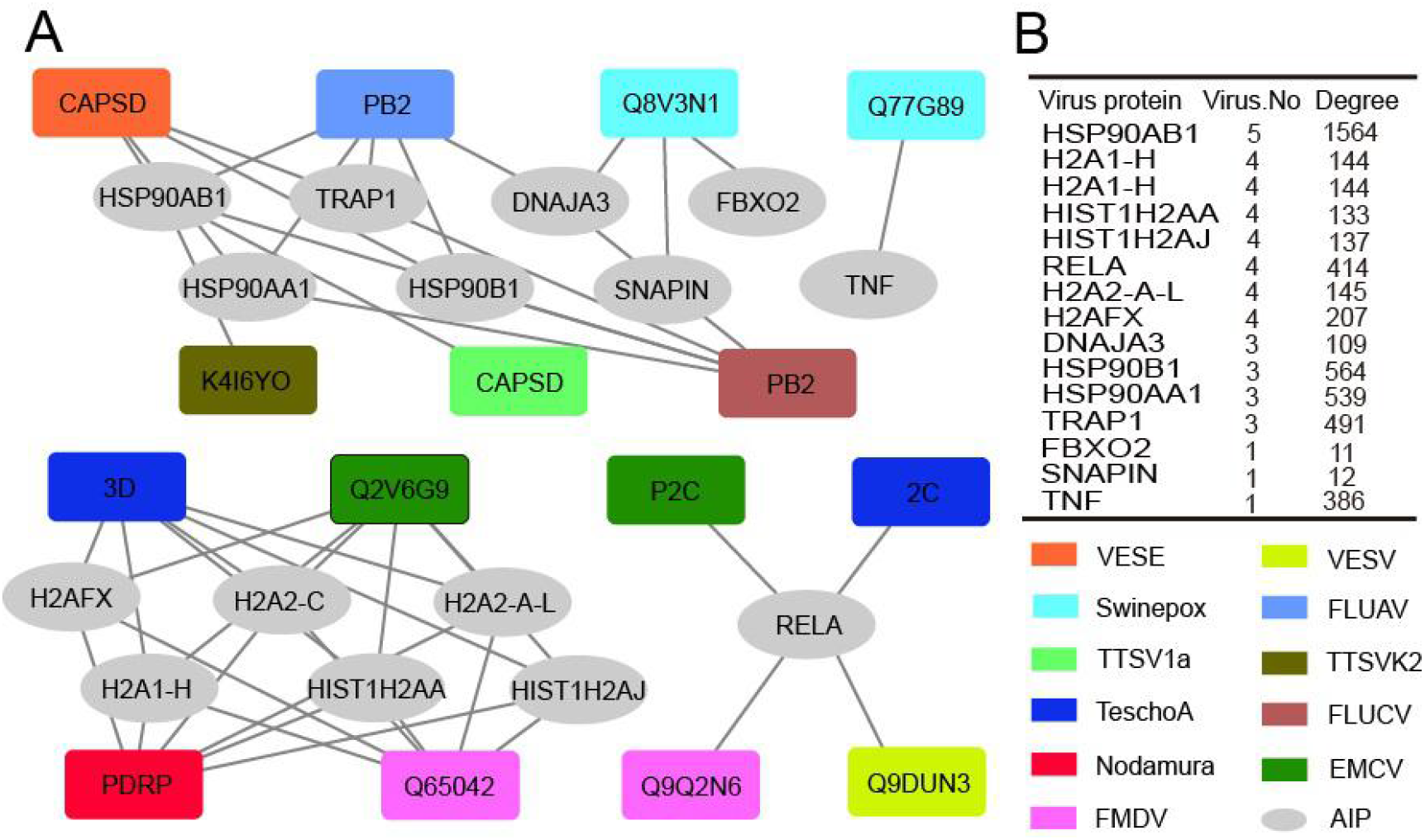
AIPs and their interactions with other viruses. (A) The PPI network between AIPs and other viruses. AIPs were represented as ellipse in gray. Viruses were represented as squares and colored according to the legend in the bottom right. VESV, Vesicular exanthema of swine virus; FLUCV, Influenza C virus; FLUAV, Influenza A virus; FMDV, Foot-and-mouth disease virus; Nodamura, Nodamura virus; EMCV, Encephalomyocarditis virus; TeschoA, Teschovirus A; Swinepox, Swinepox virus; TTSV1a, Torque teno sus virus 1a; TTSVk2, Torque teno sus virus k2; FMDV, Foot-and-mouth disease virus. (B) The number of interacted virus and the degree of AIPs in the swine PPI network. H2A1-H, histone H2A type 1-H; H2A2-C, histone H2A type 2-C; H2A2-A-L, histone H2A type 2-A-like.

### Drug prediction for treating ASFVs

The wide involvement of AIPs in PPIs between swine and multiple viruses, and the central role of AIPs in swine PPI network, suggested the possibility of their use as broad-spectrum host-dependent antiviral targets. Therefore, we attempted to predict drugs for targeting the AIPs with the help of DrugBank (Table S3). As was shown in Figure 5, a total of 167 drugs (in ellipse or square) were predicted which targeted 19 AIPs (pink circles). Most of the drugs were small molecules (colored ellipses); the other drugs were protein or peptide (colored squares). Some AIPs were targeted by multiple drugs, such as the heat shock protein 90 alpha family class A member 1 (HSP90AA1) and tumor necrosis factor (TNF). HSP90AA1 was targeted by more than 30 drugs, most of which were small molecules and were in experimental; while TNF were also targeted by more than 30 drugs, most of which were approved or investigational.

**Figure 5.**
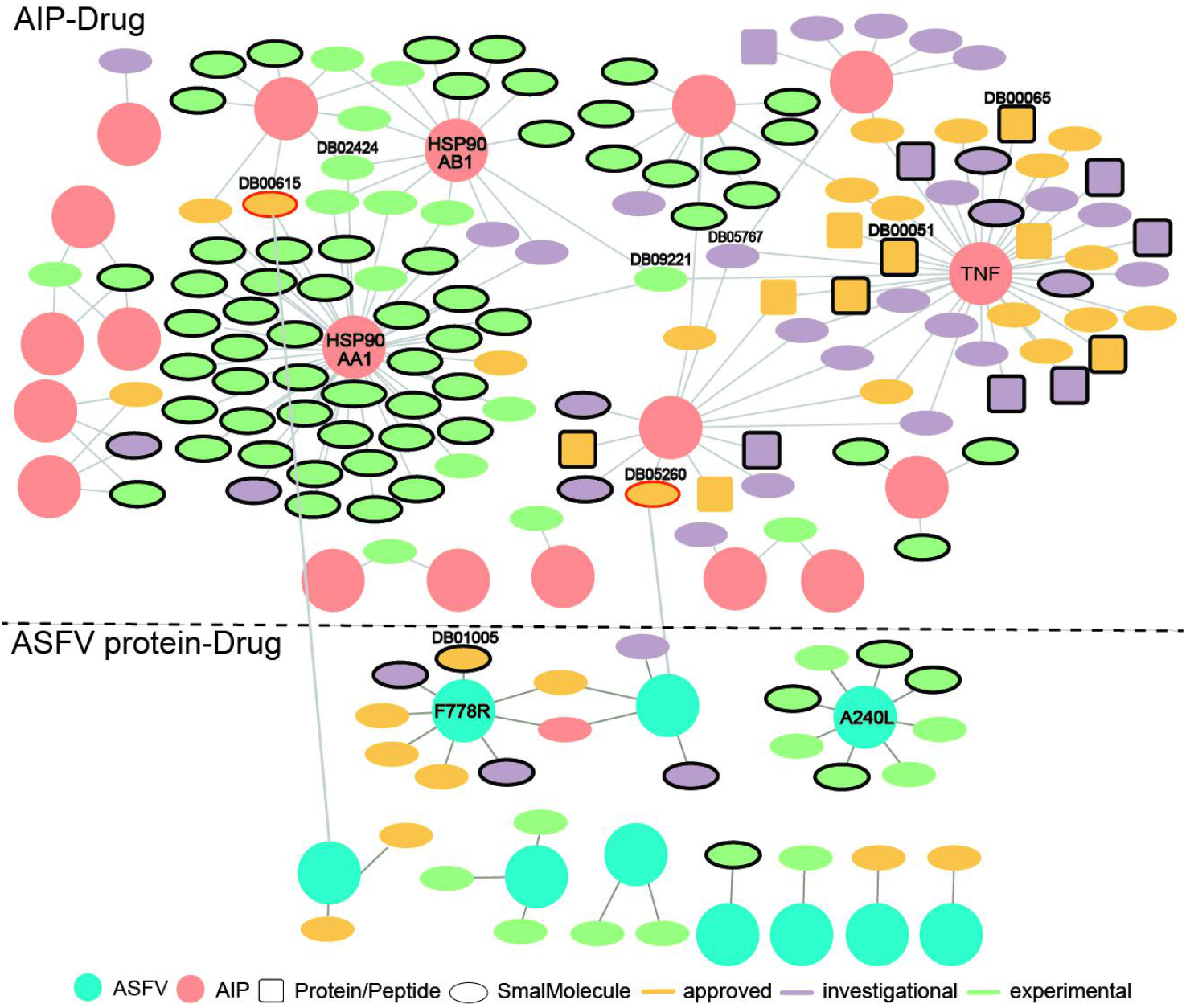
Predicted drugs targeting the AIPs and ASFV proteins. The interactions above and below the dotted line referred to those between drugs and AIPs, and those between drugs and ASFV proteins, respectively. The AIPs and ASFV proteins were represented as pink and cyan circles, respectively. Drugs of protein or peptide, and those of small molecule, were represented as squares and ellipses, respectively. Drugs in the stage of approved, investigational and experimental were colored in orange, purple and light green, respectively. Drugs which specifically targeted one AIP were highlighted in black-edge. Two drugs which targeted both the AIP and ASFV protein were highlighted in red-edge.

We also predicted drugs targeting the ASFV proteins. Twenty-nine small molecules were predicted to target ten ASFV proteins. Among them, both the proteins of F778R and A240L were targeted by eight drugs. However, these ten ASFV proteins were not involved in the PPIs between ASFV and swine. Interestingly, the Gallium nitrate (DrugBank ID: DB05260), a drug used for treating hyper-calcemia, and Rifabutin (DrugBank ID: DB00615), were observed to both target the AIPs and ASFV proteins. Both of them (highlighted in red-edge) were approved for use, suggesting their potential use for treating the ASFVs.

Some drugs were observed to have strong specificity on the ASFV protein or AIPs, such as the Hydroxyurea(DB01005), Infliximab (DB00065), Adalimumab(DB00051), and so on. Hydroxyurea specifically targeted F778R. It is an antineoplastic agent that inhibits DNA synthesis through the inhibition of ribonucleoside diphosphate reductase. It may be used to inhibit DNA synthesis of the ASFV virus, thus blocking the proliferation of the virus. Infliximab specifically targeted TNF and is primarily related to inflammation control and neurological indications. It may be used to block the necrosis during the ASFV infection.

Some drugs were observed to target multiple AIPs, such as Geldanamycin (DB02424), Polaprezinc (DB09221) and Andrographolide (DB05767). For example, Polaprezinc could target the TNF, HSP90AA1 and HSP90AB1, among which both HSP90AB1 and HSP90AA1 played a central role in the swine PPI network and swine-virus PPI network. Therefore, Polaprezinc may have a broad-spectrum effect in disrupting the swine-ASFV PPI network, and may inhibit the viral infections effectively.

## Discussion

Vaccines and antiviral-drugs are considered as the most effective tools for fighting against viruses. Unfortunately, nearly all attempts to develop vaccines against ASFVs have failed to induce effective protection ^4, 20^. Therefore, it is necessary to develop antiviral drugs against the virus. Previous studies have found several compounds which could possibly inhibit ASFV infection in vitro, including the genistein, necleoside analogues, sulfated polysaccharides, lauryl gallate, small peptide inhibitors, and so on. This study provided several candidate drugs targeting the ASFV proteins and AIPs, which may facilitate the development of more effective drugs against the virus.

In the era of systems biology, a large amount of PPIs have been accumulated, including the virus-host PPIs. This study compiled an up-to-date PPI network between ASFV and swine. Analysis of the network could help identify possible associations between viral activities and host defense strategies, which may facilitate development of potential therapies by disrupting host-virus interactions ^21^. Several AIPs and lots of AAPs were identified based on the PPI network. They were observed to interact with more proteins or have larger influences on the information flow throughout the swine PPI network than other proteins, suggesting their central roles in the swine PPI network. Some AIPs were observed to interact with multiple viruses. Therefore, the predicted drugs targeting these AIPs, such as Polaprezinc, may have a broad-spectrum effect against viral infections.

Both the AIPs and AAPs were enriched in the functions of cell death, apoptosis or necroptosis. This suggested that these processes may play an important role in viral infections. Previous studies have shown that host cells could limit ASFV replications by induction of apoptosis. For survival, ASFV encodes several anti-apoptotic proteins, such as A179L and A224L. Actually, in the late stage of ASFV infection, induction of apoptosis could favour virus spread without the activation of inflammatory responses ^22^. Therefore, the drugs which could induce or inhibit cell death, apoptosis or necroptosis, may be candidates for treatment of ASFVs. For example, the infliximab, which is a TNF blocker and primarily related to inflammation control ^23^, may be used to block the necrosis during the ASFV infection.

Lots of drugs were predicted to target the AIPs or ASFV proteins. Several strategies could be use to select the candidate drugs. For specificity, the drugs with high specificity on the AIPs or ASFV proteins could be selected, such as the Hydroxyurea and Infliximab; for broad-spectrum effect, the drugs which targeted the AIPs with high degrees in the swine PPI network, such as Polaprezinc, or those targeted multiple AIPs, such as Geldanamycin, could have a large influence on the swine PPI network or the PPIs between swine and ASFV. Two drugs, i.e., Gallium nitrate and Rifabutin, were observed to target both the AIPs and ASFV proteins. They could also be used for broad-spectrum inhibitory effect against ASFV infections.

Most antiviral drugs target the viral proteins. Drug resistance frequently appears due to rapid mutation of viruses. On the contrary, the drugs targeting the host protein may have the advantage of stable effect since the host proteins generally evolve far slower than viral proteins ^24, 25^. Besides, some host proteins may interact with multiple viruses, such as HSP90AB1 mentioned above. The drugs targeting them may have broad-spectrum antiviral effect. Bioinformatics analysis of the accumulated PPIs between virus and host cell can facilitate the identification host proteins which are vital for viral infection. As the accumulation of PPIs and the rapid development of bioinformatics methods, several antiviral molecules with reduced side effects have been proposed and validated. This study investigated the prediction of antiviral drugs against ASFV infections. The candidate drugs identified here may facilitate further development of effective drugs against the virus.

There were two limitations to this study. Firstly, the PPIs between swine and ASFV are far from complete. The ASFV encodes more than 150 proteins. Only 16 of them were involved in the PPIs analyzed here. Much more efforts are needed to generate a comprehensive PPI network between swine and ASFV. Fortunately, based on the limited PPIs between swine and ASFV, several antiviral drugs were predicted and had the potential for further development^26, 27^. Secondly, the drugs predicted here need further experimental validations. Several drugs with high specificity on AIPs, or with broad-spectrum effect, such as Polaprezinc, could be prioritized for validation.

In conclusion, this study curated a set of PPIs between swine and ASFV as more as possible and identified the AIPs which were vital for viral infection. The AIPs were observed to play a central role in swine PPI network, and also took part in interactions between swine and several other viruses. Several drugs were predicted to target the AIPs and ASFV proteins. They could be helpful for development of effective drugs against the virus.

## Acknowledgements

This work was supported by the National Key Plan for Scientific Research and Development of China (2016YFD0500300), Hunan Provincial Natural Science Foundation of China (2018JJ3039), Changsha Science and Technology Bureau (kq1801040), the National Natural Science Foundation of China (31500126 and 31671371) and the Chinese Academy of Medical Sciences (2016-I2M-1-005).

The authors have declared that no competing interests exist.

